# Ancestry-Dependent Immunologic and Prognostic Effects Characterize the Prostate Cancer Urinary Proteome

**DOI:** 10.1101/2025.08.14.670396

**Authors:** Annie Ha, Jaron Arbet, Zhuyu Qiu, Amanda Khoo, Brian P Main, Meinusha Govindarajan, Matthew Waas, Stanley K Liu, O John Semmes, Julius O Nyalwidhe, Paul C Boutros, Thomas Kislinger

**Author notes:** equal contributions (Annie Ha, Jaron Arbet, Zhuyu Qiu, Amanda Khoo).

## Abstract

Urine is an attractive biomarker analyte for non-invasive longitudinal monitoring of health and disease, particularly for diseases of the genitourinary tract, like prostate and bladder cancer. The composition of an individual’s urine reflects both genetic and lifestyle characteristics that differ across geographies and populations, like diet, hydration and other socio-economic factors. While men of African ancestry have elevated prostate cancer risk, it is unclear to what extent this influences urinary biomarkers. We therefore quantified the urinary proteomes of 329 localized prostate cancer patients: 135 self-identifying as White and 194 self-identifying as Black. We identified 110 proteins that significantly differed between these groups after controlling for age, PSA, and cISUP. Immune pathways were particularly dysregulated. The urinary proteome of Black patients harboured more features of aggressive cancers than those of grade– and PSA-matched White patients. These observations highlight the importance of controlling for race– and ancestry-associated differences in the development of urinary biomarkers.

## Introduction

Urine is an attractive source for prostate cancer (PCa) biomarker development. It contains molecules from the circulating blood that are filtered by the kidney, as well as shed and secreted from nearby urogenital organs such as bladder and prostate, which can be reflective of their physiological state^1^. Specifically, prostate is a secretory organ that can release prostatic secretion into the urine via the acini structures and urethra^2^. Hence, urine-based biomarkers can be utilized to improve current prostate cancer prognosis and diagnosis^3,4^. Several promising urine-based biomarkers have been developed to complement serum prostate-specific antigen (sPSA) for PCa diagnosis and prognosis^3,5^. However, urine composition is influenced by many factors, including time of day, hydration status, age, disease status, germline genetics and patient lifestyle ^6,7^, potentially confounding interpretation of urinary biomarkers.

The clinical outcomes of PCa are also influenced by most of these factors. Black men have been reported to have ∼67% higher incidence and ∼2-fold higher mortality rates compared to White men^8–10^. Although at least part of this divergence is confounded by socio-economic status^11,12^, recent studies show that genetic ancestry directly influences the molecular landscape of PCa, suggesting potential contributions to differences in clinical outcomes^13–15^. For instance, whole genome sequencing studies reported that tumours of African ancestry (AA) harbour higher mutational burden^13^, fewer PTEN mutations and more androgen receptor alterations compared to patients of European ancestry (EA)^15–17^. The growing evidence of ancestry-driven molecular heterogeneity in PCa suggests that ancestry-adjusted molecular biomarkers could benefit clinical management. To maximize the utility of urinary biomarkers for PCa, the influence of ancestry on the urinary proteome needs to be investigated.

To investigate the potential implications of ancestry on non-invasive markers in PCa, we assessed the associations of self-identified race and ethnicity (SIRE) on the urine proteome from a cohort of 329 individuals with localized PCa. Our study revealed proteins significantly associated with SIRE and age, independent of other clinical variables, in which immune pathways are dysregulated in Black patients. We demonstrated that the urinary proteome of Black patients more closely resembled that of biologically aggressive tumours compared to that of their White counterparts. We further reported preliminary differences in urinary marker prognosis between Black and White patients, highlighting the importance of controlling for race– and ancestry-associated differences in the development of urinary biomarkers.

## Results

### Clinical and urine proteomics characteristics of the cohort

We generated whole-proteome data on post-digital rectal exam (DRE) urine in a cohort of 329 individuals with localized PCa (**Figure 1A**, **Supplementary Figure 1A**, **Methods**). Our cohort was comprised of 135 individuals with self-identified race and ethnicity (SIRE) of White and 194 individuals of Black. Between the two groups, Black patients were slightly older (63 *vs.* 61 years, p = 1.98 x 10^-6^). Consistent with literature^18,19^, Black patients in our cohort had higher sPSA levels (6.4 *vs.* 5.0 ng/mL, p = 1.5 x 10^-7^) at diagnosis, even after adjusting for age (linear regression coefficient = 0.11, p = 1.116 x 10^-3^; **Supplementary Figures 1B-C**, **Supplementary Table 1**). The distributions of biopsy grade (cISUP grade group) and T category did not significantly differ between Black and White patients (Pearson’s X^2^ test, cISUP p = 0.06; cT p = 0.67; **Supplementary Figures 1D-E**).

**Figure 1:**
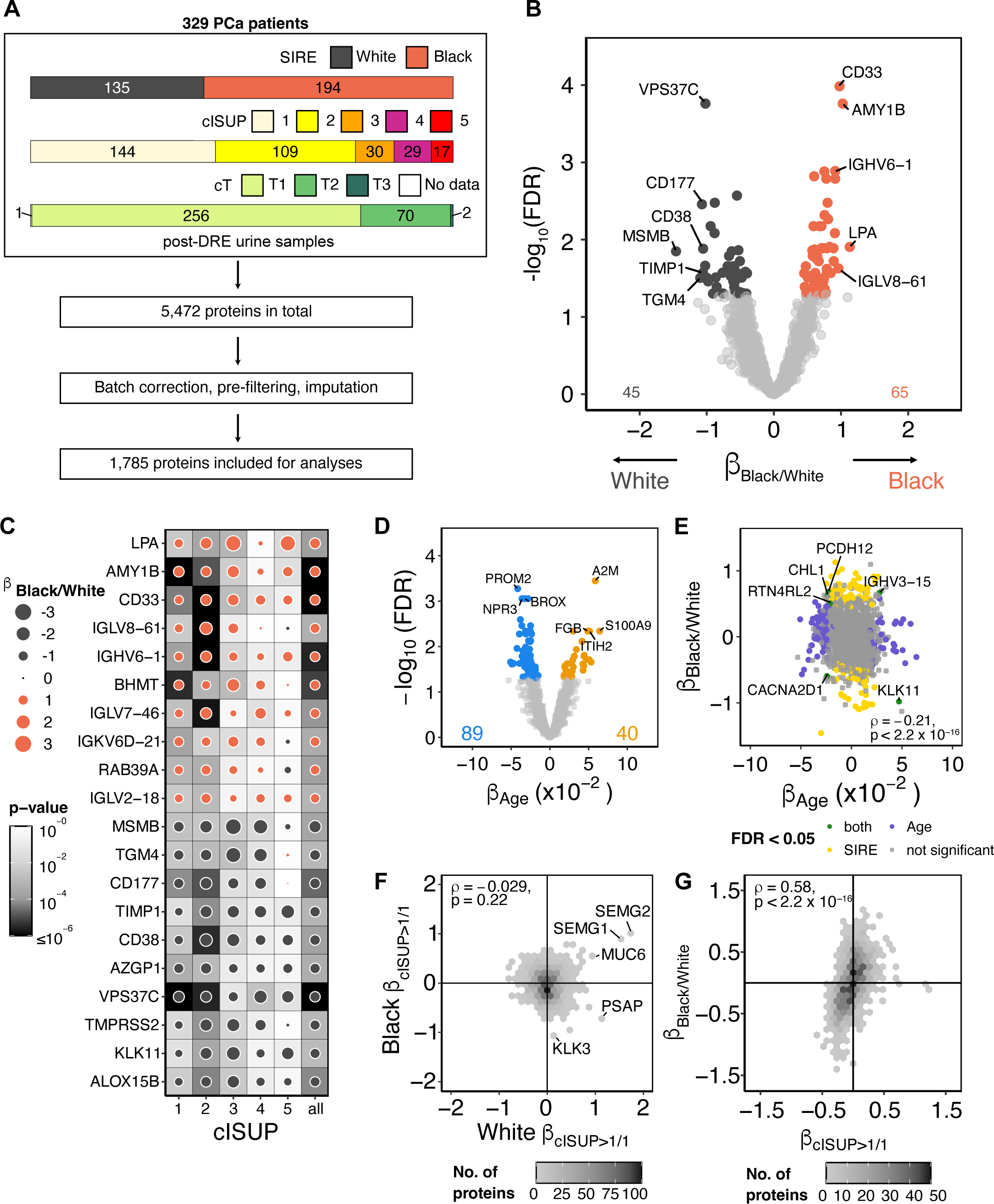
Urine proteomics heterogeneity between Black and White prostate cancer patients. (A) Overview of the urine proteomics cohort and processing workflow. (B) Volcano plot of log_2_ protein abundance association with SIRE in multivariable analysis. Regression coefficient (β_Black/White_) of proteins is shown as effect size. Proteins with significant association (FDR < 0.05) are coloured (dark grey: enriched in White patient; red orange: enriched in Black patients). (C) Dot map showing the β_Black/White_and t-statistics p-value of the top and bottom 10 proteins with the greatest effect sizes in stratified multivariable analyses within cISUP grade groups. Dot size represents the absolute effect size of each protein, background shading represents significance of the association, colour represents directionality of the association (dark grey: enriched in White patient; red orange: enriched in Black patients). (D) Volcano plot of log_2_ protein abundance association with age in multivariable analysis. Significantly associated proteins are coloured (blue: negatively associated; orange: positively associated) (E) Spearman’s correlation of proteins associations for SIRE and Age. Significantly associated proteins (FDR < 0.05) are coloured (green: significant in both variables; purple: significant in age only; yellow: significant in SIRE only; grey: not significant). (F – G) Spearman’s correlations of log_2_ protein abundance associations with (F) disease stage (cISUP > 1/cISUP1) between Black and White patients from stratified multivariable analyses within SIRE, (G) SIRE and disease stage from multivariable analysis not stratified.

Our urine proteomics pipeline detected 5,472 proteins in total across all patients including various prostate-specific proteins. In our cohort, White patients’ urine samples yielded slightly more proteins compared to Black patients (**Supplementary Figure 2A**). To assess general variance of the urinary proteome, principal component analysis revealed no clear clustering patterns prior to (**Supplementary Figure 2B**) and after data normalization (**Supplementary Figure 2C**). The generated proteomics data spans ∼6 orders of magnitude and detected various prostate-derived proteins such as KLK3, ACP3, TGM4, and STEAP2 (**Supplementary Figure 2D**). Following normalization of proteomics data, downstream analyses focused on the 1,785 consistently detected proteins (observed in ≥50% of patients in each SIRE group) (**Supplementary Figure 2E**). Between SIRE groups, there were no proteins highly detected (>80%) in one group only (**Supplementary Table 2**).

### Race and ethnicity urine proteome associations

As the molecular landscape of PCa is influenced by variables such as age and tumour grade^6,14,20^, a multivariable analysis adjusting for cISUP grade group^21^, age, and sPSA level was used to quantify associations of urinary protein abundance with SIRE. We identified 110 proteins significantly associated with SIRE (**Figure 1B**, **Supplementary Table 3**), with top-ranking proteins showing consistent SIRE-driven protein abundance changes across tumour grade groups (**Figure 1C**, **Supplementary Figure 3A-B**). Immune-related proteins (*e.g*. immunoglobulins) were more abundant in Black patient urines, while prostate-derived proteins (*e.g.* MSMB, TGM4, TMPRSS2) were more abundant in White patient urines. Multivariable analyses also uncovered 129 proteins significantly associated with age (**Figure 1D**), and that proteins significantly associated with both variables are likely independent (Hypergeometric test; p = 0.82; **Figure 1E**). These data show that the urinary proteome is heavily influenced by SIRE, even after controlling for age and clinical variables (**Supplementary Figure 3C**).

To examine the correlation of urinary protein abundance associations between SIRE and tumour aggressiveness, we compared the effect sizes between cISUP 1 (*i.e.,* clinically insignificant) and cISUP > 1 tumours (*i.e.,* clinically significant). To compare grade-group differences between SIRE, protein abundance associations with grade-group is assessed using multivariable analysis within Black and White patients. Black and White patients show fundamentally different proteins associated with tumour grade in this stratified analysis (**Figure 1F**). For example, while SEMG1 and SEMG2 are associated with clinically significant tumours in both groups, most proteins are only associated in one group or the other. Even PSA itself (KLK3) is only associated with clinically significant tumours in White patients. On the other hand, to assess the correlation between SIRE and tumour grade, we performed multivariable analysis assessing protein abundance associations with tumour grade adjusting for SIRE, age, and sPSA. Within the same model, we observed that proteins associated with SIRE were preferentially associated with disease grade (**Figure 1G**). The observations suggest that the urinary proteomes of Black patients are different from grade– and age-matched White patients, and resembles features of higher-grade disease even when diagnosed with lower-grade disease.

### Integrative-omics analysis revealed concordant ancestry and SIRE associations

To assess whether the urine proteomics observations were generalizable, we replicated our analysis in the TCGA tissue transcriptomics data, using genetically determined ancestry^22^ (**Figure 2A**). Concordant with urinary proteomics clustering patterns, principal component analysis showed no clear pattern separated by ancestry (**Supplementary Figure 3D**). After controlling for age, sPSA and cISUP, 829 genes were significantly associated with ancestry (**Figure 2B**, **Supplementary Table 4**). Grade-group differences were moderately correlated between ancestries (**Supplementary Figure 3E**), and the association of ancestry-differences with grade observed in the urine was not replicated in tissue transcriptomes (**Supplementary Figure 3F**). While direct comparisons between the-omics modalities are confounded by various levels of regulations (*e.g.* complex post-translational modifications and transport mechanisms), we observed many proteins associated with SIRE or ancestry in both urine and tissue (**Figure 2C**). These included elevation of prostate-derived proteins in White patient urine and tissue (*e.g.* TGM4, MSMB, CD177) and elevation of blood-derived proteins (*e.g.* CETP, LPA) in Black patient urine and tissue (**Figure 2D**).

**Figure 2:**
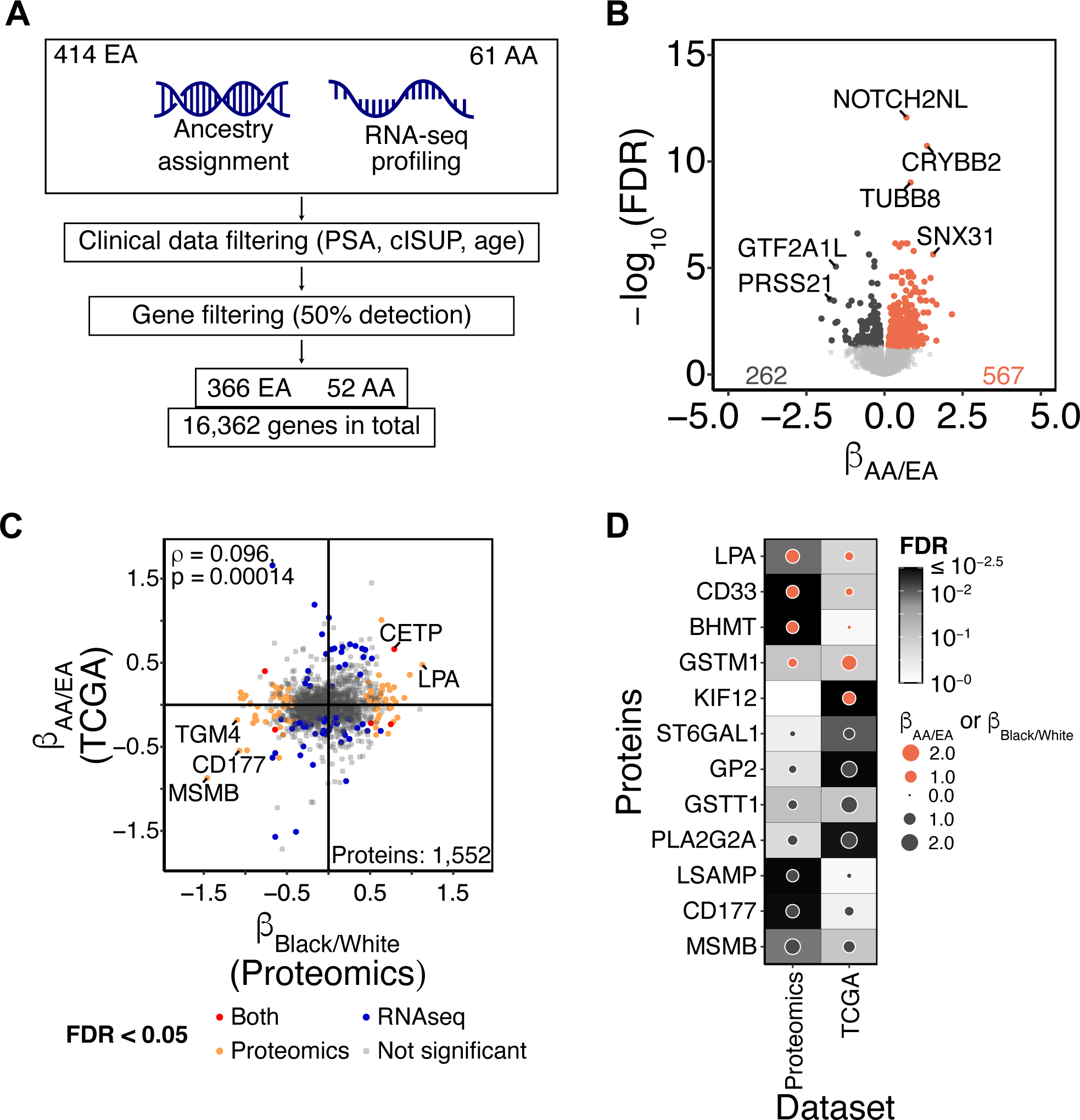
Verification of ancestry-driven differences in TCGA tissue transcriptomics. (A) Overall schematic of the TCGA PRAD transcriptomics dataset filtering criteria. Patients with missing sPSA, cISUP or age information are removed, genes detected in at least 50% of both ancestry groups are included (African Ancestry: AA; European Ancestry: EA). (B) Volcano plot of the log_2_RSEM gene expression associated with genomically inferred ancestry^22^ (African Ancestry: AA; European Ancestry: EA) in multivariable analysis. Regression coefficient (β_AA/EA_) of proteins is shown as effect size. Proteins with significant association (FDR < 0.05) are coloured (dark grey: enriched in EA patients; red orange: enriched in AA patients). (C) Spearman correlations of the tissue transcriptomics β_AA/EA_ and urine proteomics β_Black/White_. Proteins with significant associations are coloured (FDR < 0.05). The colour represents statistical significance in the independent analyses (Red: significant in both datasets; yellow: significant in urine proteomics only; blue: significant in RNA-Seq only; grey: not significant). (D) Dot map showing proteins and genes with same directionality in association with ancestry or SIRE in independent multivariable analysis within each dataset. Dot size represents the absolute effect size of each protein from independent dataset, shading represents significance of the association, colour represents directionality of the association (dark grey: enriched in EA/White patients; red orange: enriched in AA/Black patients).

### Inflammation and immune pathways enriched in Black patients

Next, we sought to quantify whether the urinary proteome differences reflected specific biological pathways. To investigate the inter-individual heterogeneity of pathway enrichment patterns, we performed sample-wise gene set variation analysis (GSVA)^23^. Unsupervised hierarchical clustering of sample-wise gene set enrichment scores revealed two distinct clusters predominantly separated by SIRE (Fisher’s Exact test, p = 2.764 x 10^-6^; **Figure 3A**, **Supplementary Figure 4A**), with immune pathways enriched in Black patients and cell cycle and vesicle-mediated transport pathways enriched in White patients (**Supplementary Table 3**). In particular, immunoglobulin production is significantly enriched in Black patients’ urine samples (**Figure 3B**), and androgen response is significantly enriched in White patients’ urine samples (**Figure 3C**).

**Figure 3:**
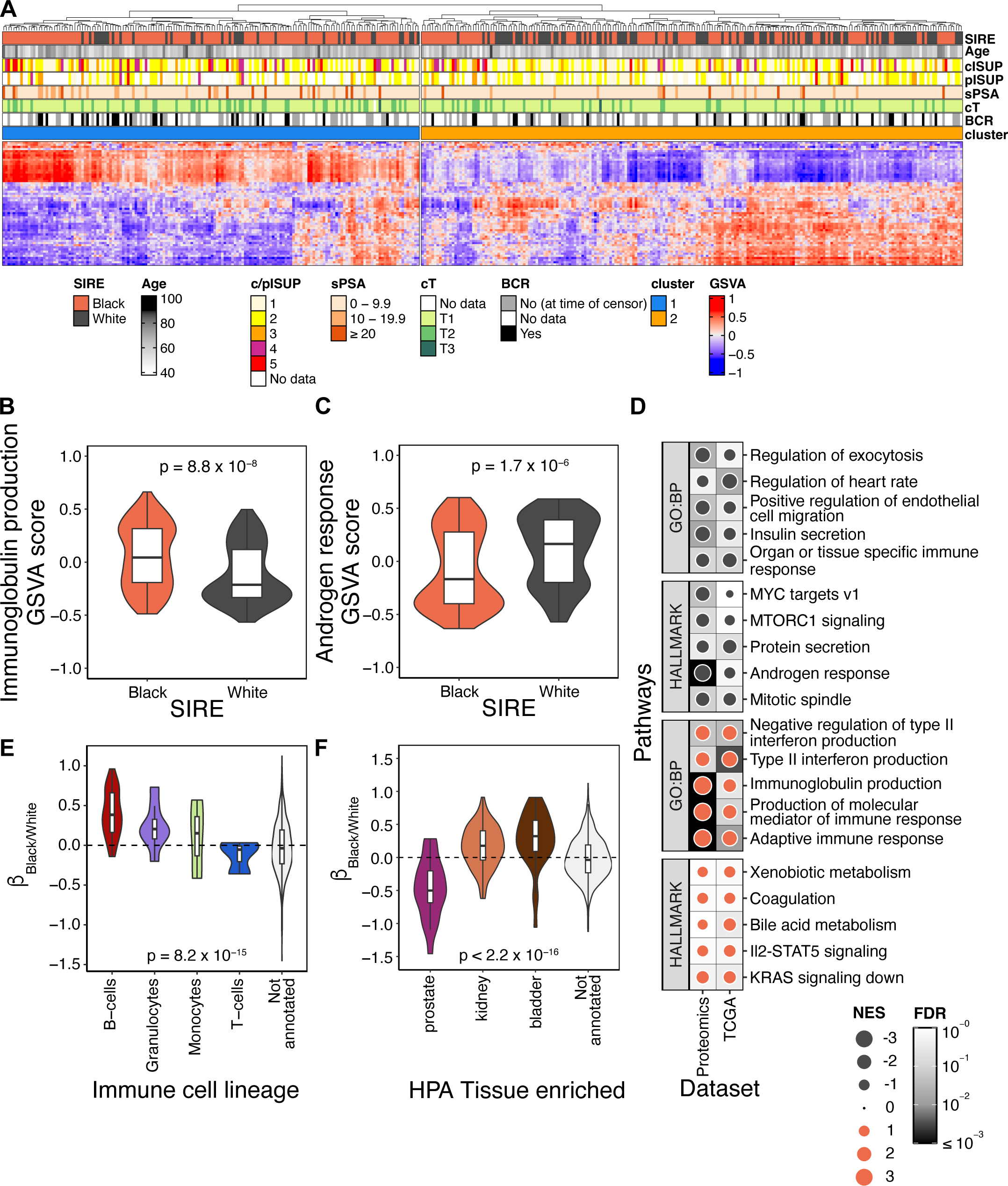
Immune-related pathways enriched in Black patients. (A) Heatmap of GSVA scores of REACTOME pathways. Pathways with GSVA scores significantly associated (FDR < 0.1) with SIRE in multivariable analysis are included in this heatmap. Heatmap is split into two by the second degree of column dendrogram from unsupervised hierarchical clustering. Column annotations represent clinical characteristics of the patient samples. Row names are listed in **Supplementary Table 3**. (B – C) Violin plots and boxplots of the GSVA score for (B) GO:BP immunoglobulin production pathway and (C) HALLMARK androgen response pathway. P-values from Mann-Whiteney U test between Black and White patients are shown. (D) Dot map showing enriched pathways from pre-ranked GSEA analysis. Pathways with same directionality for both urine proteomics and tissue transcriptomics datasets are shown. Dot size represents absolute normalized enrichment score (NES), colour represents directionality (dark grey: enriched in White/EA patients; red orange: enriched in Black/AA patients), and the background shading represents the adjusted p-value (FDR < 0.25). (E – F) Violin plot and boxplot of the β_Black/White_annotated against the Human Protein Atlas (E) immune cell lineage and (F) tissue enriched dataset. P-value from ANOVA is shown. P-values from t-test for each annotation against 0 are listed in **Supplementary Table 3**.

To examine cohort-level pathway enrichment trends, we performed global pathway enrichment analyses (GSEA)^24^. Global pathway analysis revealed consistent enrichment terms in TCGA RNA-seq data and our urinary protein data, including the increased expression of immune pathways in Black patients and androgen response pathways in White patients (**Figure 3D**). To further characterize the proteins associated with the significantly enriched pathways, we compared the abundance of proteins involved in various immune cell lineages and tissue origins. B-cell associated proteins were relatively enriched in Black patients (**Figure 3E**), whereas prostate-specific and extracellular vesicle associated proteins were relatively depleted in Black patients (**Figure 3F**, **Supplementary Figure 4B**, **Supplementary Table 3**). While prostate-specific proteins were depleted in Black patient urine, they were also depleted in urine from patients with elevated disease burden^25^ (**Supplementary Figure 4C**), further supporting the association between the urinary proteome of Black patients and tumours of more aggressive stages, even when diagnosed with lower grade tumours. The concordant pathway-level observations between urine and tissue (**Supplementary Figure 4D**) support the idea that ancestry is strongly influencing disease molecular heterogeneity.

### Urinary protein prognosis differs by race and ethnicity

To evaluate whether proteins prognostic for biochemical recurrence (BCR) vary by SIRE, we used multivariable Cox proportional hazards modeling. To limit clinical heterogeneity, we analyzed only the 133 intermediate risk patients (cISUP 2 – 3, sPSA 10 – 20 ng/mL) who received definitive treatment ^26^ (**Supplementary Table 5**). Overall BCR-free survival was not associated with age and SIRE (**Figure 4A-B**). To assess whether prognosis by protein abundance depends on SIRE, we examined the interaction of these variables. There is evidence indicating differences in the prognostic potential of proteins based on SIRE, though statistical significance was not achieved after multiple testing correction (minimum FDR = 0.18; **Figure 4C**), possibly due to limited sample size. Despite this, several proteins demonstrated opposing trends of BCR-free survival between Black and White patients (**Supplementary Figure 5A**). For instance, the overall CTSG abundance was not associated with outcome (**Figure 4D**) but was associated after adjustment for SIRE in our model (HR_interaction_: 0.36, FDR = 0.18). Low CTSG abundance was found to be significantly associated with worse outcome in Black patients, but better outcome in White patients (**Figures 4E – F**). A similar trend was also observed for KRT14 in our dataset (**Supplementary Figures 5B – D**). Such discordance suggests that prognostic urinary biomarkers may have varying efficacies between Black and White patients, though larger cohorts are needed to verify these observed results given our limited power.

**Figure 4:**
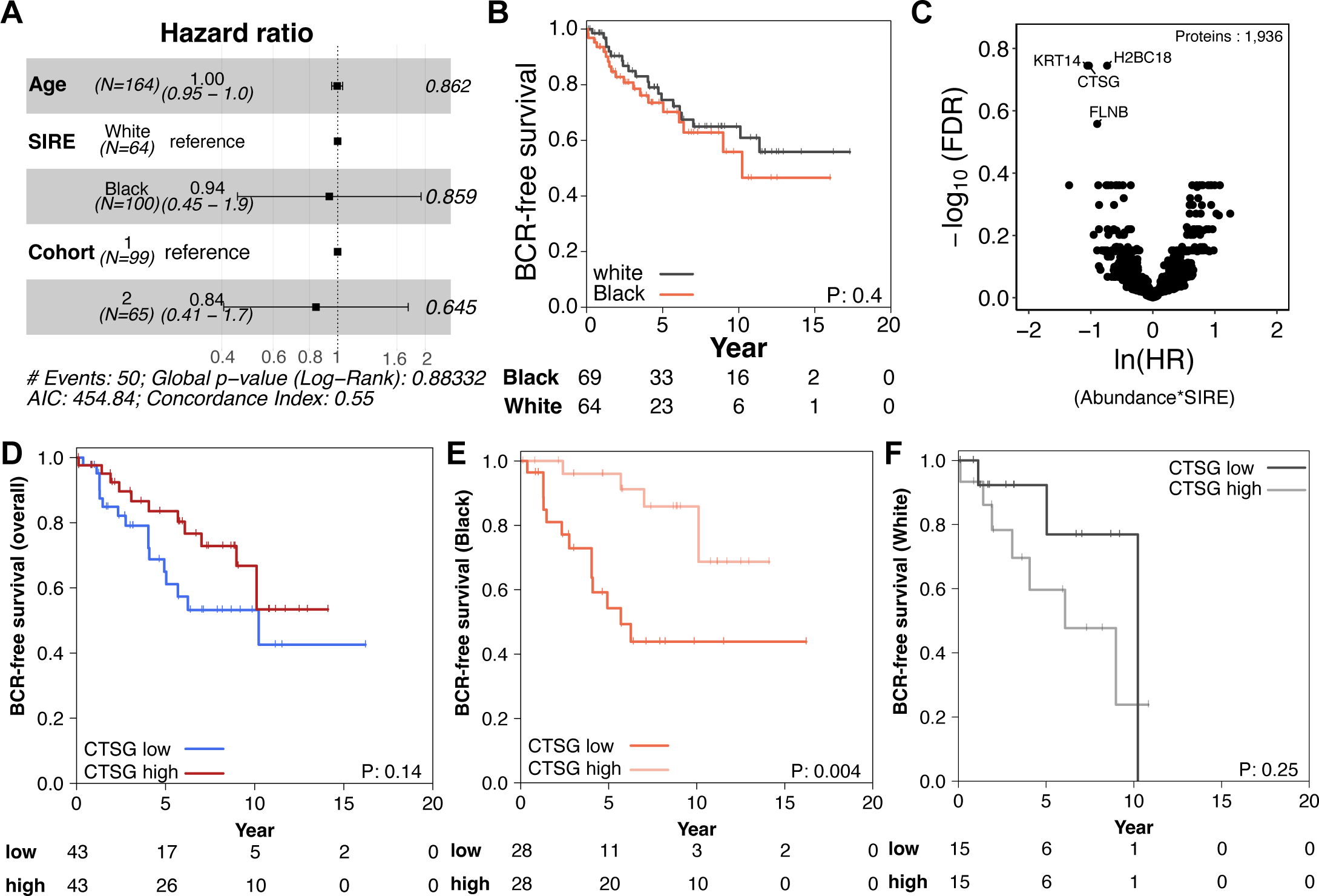
Urinary prognostic markers differ by SIRE. (A) Forest plot of hazard ratios (HR) for each clinical term for biochemical recurrence (BCR)-free survival in a multivariable Cox-proportional hazards (Cox-PH) model for intermediate risk patients. Box represents estimated HRs, whiskers represent the 95% confidence interval of HR. Wald-test p-values for each covariate is shown on the right. (B) Kaplan-Meier (KM) plot of overall biochemical recurrence (BCR)-free survival in intermediate risk patients stratified by SIRE. P-value from log-rank test is shown. (C) Volcano plot of the interaction between SIRE and protein abundance association with BCR. Regression coefficient of the interaction term is displayed as ln(HR). P-values are corrected by BH-FDR method. (D) Kaplan-Meier curve of biochemical recurrence (BCR)-free survival in intermediate risk patients stratified by overall urinary CTSG abundance (median dichotomized). P-value from log-rank test is shown. (Blue: CTSG low; red: CTSG high) (E-F) Kaplan-Meier curve of BCR-free survival in intermediate risk (E) White and (F) Black patients stratified by CTSG urinary protein abundance level (median dichotomization). P-values are from the log-rank test.

## Discussion

Many urine-based biomarkers are actively being investigated to complement sPSA for PCa diagnosis and prognosis^1,3,5,27^. While the urinary proteome varies with disease stage^6,28^, it can also be influenced by various lifestyle factors driven by ancestry. Growing evidence demonstrating tumour molecular heterogeneity driven by ancestry also suggests potential implications in the currently established diagnostic and prognostic protocols, leading to inaccuracies in staging of the disease^13–15,17,22,29,30^. To improve overall outcomes of PCa in diverse populations, influences of ancestry on non-invasive markers are needed. Here, we report the first and the largest study investigating the ancestry– and ethnicity-driven influences on the urinary proteome of patients with localized PCa.

We generated whole-proteome profiles from urine samples of 329 patients with localized PCa and characterized the differences between self-identified Black and White patients. Our analyses revealed that even in patients with matched disease grade, both age and SIRE are associated with hundreds of urinary features, independently. In particular, 110 proteins (*i.e.* ∼10% of the modeled proteome) were significantly associated with SIRE, independent of other clinical variables. These observed differences are enriched in specific biological pathways such as immune and inflammatory responses and androgen response. Our analyses also demonstrated that the urinary proteome of Black patients resembled more closely to that of higher-grade disease than that of grade– and PSA-matched White patients. Consistent with other studies, immune dysregulation and upregulation of immune markers have been observed in the prostate tumours of Black patients^22,31–34^. While we are able to capture the enrichment of immune signatures in the urinary proteome, the growing evidence of ancestry-driven heterogeneity highlights the enrichment of immunological landscapes in the tumour of Black patients, which may render varying responses to immunotherapies and serve as new avenues for targeted therapies^35,36^.

Our analyses also revealed several proteins with inverse prognostic effects between Black and White patients. Although sample size is limited in the survival analysis for biochemical recurrence association, interaction effects between protein abundance and SIRE were observed. Specifically, we highlighted the examples of urinary CTSG and KRT14 prognosis depends on SIRE. That is, both proteins were not associated with BCR when considered across all patients, but were associated with outcome once stratified within SIRE groups. Furthermore, we also observed that KLK3 (*i.e.* PSA) is negatively associated with tumour grade in the urine of Black patients, which is the opposite effect in White patients. The reported discordance in non-invasive markers between Black and White patient suggests that currently established biomarkers may have varying performance across diverse ancestry and ethnicity populations^37^. Hence, identifying ancestry-or race and ethnicity-adjusted molecular biomarkers may benefit overall diagnostic accuracy, and our findings emphasize the need of larger diverse cohorts for the development of clinically-generalizable biomarkers.

The findings in this study provide an in-depth insight into the differences of the ancestry-driven urinary proteome, though several limitations remain. Due to limited access of matched tumour samples, genetically inferred ancestry information was unavailable, although high correlations between genetic ancestry and SIRE have been previously reported^22,38–40^. On the other hand, while we reported that urinary markers have varying prognostic potentials based on self-identified race and ethnicity, limited sample size and events (BCR) challenged statistical significance. Larger urine cohorts consisting of patients managed by definitive treatment with long-term follow-up (∼15 years) are needed for validation of our observed results. However, such cohorts are currently unavailable within our group, and will require additional biobanking or integration with other clinical centers, both associated with significant lead time, representing a limitation of the study. While our analyses considered other clinical variables such as tumour grade and age, we were unable to account for residual confounding factors from other socio-economic and lifestyle variables (*e.g.* income, education, body-mass-index, diet) as they were unavailable.

In summary, we reported the ancestry-driven heterogeneity of the urinary proteome of patients with localized PCa. Multivariable analyses revealed 110 proteins independently associated with self-identified race and ethnicity, with inflammatory and immune responses significantly enriched in Black patients. Stratified analyses also revealed that the urinary proteome of Black patients harboured more features of aggressive cancer than that of their White counterparts. Survival analyses revealed inverse prognostic effects of urinary markers between Black and White patients. Hence, the growing evidence of tumour heterogeneity driven by ancestry and lifestyle should be critically considered in biomarker development. Further validation with larger and even more heterogeneous cohorts are needed to better understand tumour biology for improved precision medicine, as well as to identify more clinically-generalizable biomarkers.

## Online Methods

### Patient cohort

Clinical samples were obtained from men following informed consent and use of Institutional Review Board approved protocols at the Eastern Virginia Medical School (EVMS, Norfolk, Virginia, USA, IRT# 06-12-FB-0343) and the Research Ethics Review Board at the University Health Network (UHN, Toronto, Ontario, Canada, 10-0159 and 19-5009). The discovery cohort included patients diagnosed with localized prostate cancer from 1990 to 2017. Patients with *de novo* metastasis, benign prostatic hyperplasia, self-identified race and ethnicity (SIRE) groups other than Black or White, and those with missing clinical biopsy information were excluded from subsequent SIRE comparisons, detailed exclusion criteria are listed in **Supplementary Figure 1**. Longitudinal urine samples or post-treatment samples were also excluded from this study. Detailed clinical information is listed in **Supplementary Table 1**.

### Urine collection

First catch urine (15 mL) was collected post digital rectal exam as previously described^6,41^. Urine was centrifuged at 2000 x *g* for 15 minutes at 4°C to remove cellular debris and the supernatant was stored at –80°C.

### Urine proteomics

Samples were randomized to random numbers generated by uniform distribution prior to processing. Urines were prepared for proteomics using the MStern protocol as previously described^6,41,42^. For each sample, 2 pmol of *Saccharomyces cerevisiae invertase* (SUC2) (Sigma-Aldrich) was added as a sample processing control. Proteins were reduced with 5 mM dithiothreitol and incubated for 30 min at 60°C. Proteins were then alkylated with 25 mM iodoacetamide at room temperature for 30 min in the dark. The MStern membrane (Millipore Sigma, MSIP4510) was equilibrated with 50 µL of 70% ethanol and washed twice with 100 mM ammonium bicarbonate (ABC). 250 µL of urine was added to each well and passed through the membrane by vacuum suction. Proteins were washed twice with 100 µL ABC. Proteins were digested with 1 µg of mass spectrometry grade Trypsin/Lys-C enzyme mix (Promega) in 50 µL of digestion buffer (100 mM ABC, 1 mM CaCl_2_, 5% acetonitrile [ACN]). The digestion buffer was passed through the membrane by centrifugation with a collection plate, and the flow through was reapplied on top of the membrane. Proteins were then digested at 37°C for 4 hours. Peptides were collected by centrifugation. Remaining bound peptides were eluted with 50 µL ACN twice. Samples were dried in a SpeedVac vacuum concentrator (Thermo) and resuspended in 0.1% trifluoroacetic acid in water. Peptides were desalted using in-house solid phase extraction stage tips with 3 plugs of 3M^TM^ Empore^TM^ C18 membrane. Desalted peptides were dried in SpeedVac vacuum concentrator and reconstituted in 0.1% formic acid (FA) in water. Peptide concentration was determined by NanoDrop (Thermo). Synthetic iRT peptides ^43^ (Biognosys) were resuspended in 50 µL of Dissolution Buffer as per manufacturer’s instructions and spiked into each sample at a 1:10 (v/v) iRT to peptide ratio. For each sample, 2 µg of peptides were used for liquid chromatography-mass spectrometry (LC-MS) as previously described^41^.

### Liquid Chromatography

Peptides were separated by reversed phase chromatography on an EASY-nanoLC 1000 (Thermo), equipped with a 1.5 cm trap column (1.5 cm x 75 µm) and an EASY-Spray reversed-phase HPLC analytical column (500 mm x 75 µm inner diameter x 2 µm, C18 beads). Solvent A (0.1% FA in water) and solvent B (0.1% FA in ACN) were used. For samples acquired by DDA-MS, peptides were eluted with a non-linear gradient of 130 min, starting with 5% B for 5 min, 5% to 25% B over 100 min, then from 25% to 48%B in 10 minutes, followed by a wash step of 48% to 95% B for 4 minutes and 95% B for 10 minutes. For samples acquired by DIA-MS, peptides were eluted with a 50 minute non-linear gradient, starting with 0% to 8% B for 5 minutes, 8% to 24% B for 30 minutes, 24% to 48% B for 5 minutes, 48% to 95% B for 5 minutes, followed by a wash step of 95% B for 5 minutes.

### DDA-MS acquisition

To generate a spectral library for this cohort, all samples were acquired with DDA-MS on a Q-Exactive HF mass spectrometer (Thermo). MS1 survey scans were acquired with a mass range from 350 – 1800 *m/z*, at 60,000 resolution, 40 ms maximum injection time (maxIT), and automatic gain control (AGC) 3 x 10^6^. Top 15 mode was selected for MS2 fragmentation using high energy collision dissociation at normalised collision energy 27, maxIT of 55 ms and dynamic exclusion of 40 s.

### DIA-MS acquisition

DIA-MS data were acquired on a Q-Exactive HF mass spectrometer for the urine proteomics cohort. MS1 full survey scan was acquired from 350 – 1800 *m/z* with resolution of 60,000, 40 ms maxIT, with AGC 3 x 10^6^. MS2 scans were acquired with isolation width of 16 *m/z* from 400 – 1,000 *m/z* mass range with loop count of 38 (30,000 resolution, 55 ms maxIT, and AGC 1 x 10^6^). Isolation windows were offset by 8 *m/z* per cycle^44^, window placements were optimized by Skyline^45^.

### Spectral library generation

The full urine proteome spectral library was generated using FragPipe^46^ (v.20.0) with workflow “DIA_SpecLib_Quant” using all DDA-MS files. The platform was configured with MSFragger^47^ (v.3.8), Philosopher^48^ (v.5.0.0), EasyPQP (v.0.1.40, https://github.com/grosenberger/easypqp), DIA-NN^49^ (v.1.8.2), and IonQuant^50^ (v.1.9.8). DDA-MS.raw files were imported to the workflow and searched against the human UniProt-SwissProt database (2023-07-27, 42,450 sequences from canonical and isoform) using MSFragger. Database search was performed with the default settings. Search results were statistically validated through Philosopher and MSBooster^51^, peptide spectrum matches with FDR < 1% were used for spectral library generation. Retention time (RT) alignment and consensus spectra generation were performed by EasyPQP (RT loess fraction = 0.05, CiRT was chosen for RT alignment). DIA-NN was turned off for library generation. The output library.tsv was used for DIA-MS data analysis.

### DIA-MS data processing

DIA-MS.raw files were converted to.mzML files using ProteoWizard MSConvert (v.3.0). The converted DIA-MS files were then searched against the urine spectral library with FragPipe. Spectral library search and peptide quantification were performed with DIA-NN (v.1.8.2) with “unrelated runs” checked, using the high precision “AnyLC” algorithm. Precursors and proteins with FDR < 1% were reported in the output tables diann-output.tsv files individually. Gene-level MaxLFQ intensity reported was used for downstream analysis for protein groups with FDR < 1%.

### Proteomics data normalization and imputation

R package limma (v.3.56.2)^52^ was used for normalization and batch correction of the proteomics data. MaxLFQ intensities across runs were normalized using the normalizeCyclicLoess function for all protein groups detected. The removeBatchEffect function was used to correct batch effects between the two cohorts, stratified within SIRE. Variance between samples was assessed using principal component analysis (PCA) and the limma plotMDS function, both prior and post-normalization and correction. Proteins that were detected in at least 50% in both Black and White patients were retained for further analysis (**Supplementary Table 2**). Missing values were imputed using the R package missForest (v.1.5)^53^ resampling without replacement. To evaluate data quality after normalization, batch correction and imputation, we compared the differences of 48 matched samples processed in the two batches using two-sided paired Mann-Whitney U test. Eleven proteins had FDR < 0.05^54^, and were removed from further analysis. No further data normalization or correction was applied to the data to preserve biological variance.

### Urine differential protein abundance analysis

A univariate analysis was performed to compare each protein between Black and White patients. The log_2_ fold change of each protein was calculated using the median protein abundance. P-values were calculated from a two-sided, unpaired Mann-Whitney U-test with the wilcox.test function in R. In addition, multivariable linear regression was used to calculate the association between SIRE groups and protein abundance while adjusting for batch and clinical variables. Clinical variables considered include clinical International Society of Urological Pathology (cISUP) Grade Group treated as categorical covariate, serum PSA (sPSA, ng/mL) and age at diagnosis treated as continuous covariates (**Supplementary Table 3**). To assess protein differences within tumour grade, we stratified the multivariable linear regression by cISUP Grade Group, omitting cISUP as a covariate. To assess protein abundance association between clinically significant (cISUP >1) and clinically insignificant (cISUP 1) tumours, cISUP was modified into cISUP > 1 (2-5) and cISUP 1 groups from the multivariable linear regression. P-values were adjusted for multiple-testing using the Benjamini-Hochberg method^54^. Collinearity between covariates was assessed using the variance inflation factor function from the MASS (v.7.3-60.0.1) R package^55^.

### Tissue differential RNA abundance analysis

Prostate cancer tissue RNA-Seq data from The Cancer Genome Atlas (TCGA) were downloaded from UCSC Xena (xena.ucsc.edu, dataset ID TCGA.PRAD.sampleMap/HiSeqV2)^56^. Patients (n = 54) with missing sPSA values or cISUP Grade Group at diagnosis were removed from subsequent analyses. Genes detected in at least 50% of included samples (n = 209, log_2_RSEM > 0) were included. A multivariable linear regression model was used to estimate the association between genetically inferred ancestry assignment (European Ancestry: EA, African Ancestry: AA)^22,38,57^ and gene expression (log_2_ RSEM) while adjusting for age, sPSA and cISUP Grade Group. P-values were adjusted for multiple testing as mentioned above^54^.

### Cell Type Specificity Analysis

Bladder, kidney and prostate enriched genes were retrieved from the Human Protein Atlas (HPA) tissue enriched section (n_Bladder_ = 191, n_Kidney_ = 460, n_Prostate_ = 126). Genes that were enriched in more than one tissue were removed from analysis. Immune lineage annotations for B-cells, T-cells, NK-cells, Monocytes, Granulocytes and Dendritic cells were retrieved from the HPA Immune Cell section^60^. Proteins enriched in prostate urinary extracellular vesicles (uEV) or not (urine soluble proteins [uSP]) were retrieved Khoo *et al.* ^6^ (n_uEV-P20_ = 248, n_uEV-P150_ = 248, n_uSP_ = 184).

### Pathway Enrichment Analysis

Pre-ranked gene set enrichment analysis was performed using GSEA (v.4.3.3)^24^. Proteins included in the “**Urine differential protein abundance analysis”** section were ranked by the regression coefficient for SIRE against protein abundance. Genes included in the “**Tissue differential gene expression analysis”** section were ranked by the regression coefficient for ancestry against gene expression. The ranked list was queried against Hallmark^61^, REACTOME^62^ and Gene Ontology: Biological Process(GO:BP)^63^ databases. Pathways with FDR < 25% were kept for further analysis, and were visualized in Cytoscape (v.3.9.1)^64^ using the EnrichmentMap (v.3.3.4), annotated with the AutoAnnotate (v.1.3.5) and clusterMaker App (v.2.2).

### Gene Set Variation Analysis

Gene set variation analysis (GSVA, v.1.48.3, default settings)^23^ was used to calculate a univariate pathway enrichment score for each pathway in the GO:BP, HALLMARK, KEGG and REACTOME databases. Proteins included in the “**Urine differential protein abundance analysis**” section were used. Gene sets with minimum and maximum sizes of 15 and 200 were selected. The GSVA scores for proteomics data were tested for association with SIRE group with from the same multivariable model as mentioned in the “**Urine differential protein abundance analysis”** section. GSVA scores from REACTOME pathways that have association with SIRE (FDR < 0.1) from the multivariable linear model were visualized in a heatmap using ComplexHeatmap (v.2.16.0). P-values reported from the multivariable model were adjusted for multiple testing as described above^54^.

### Time-To-Event Analyses

Patients of intermediate risk group defined by the D’Amico risk stratification system (cISUP 2 – 3, sPSA 10-20 ng/mL, clinical T2b)^65^ and those who received definitive treatments (radical prostatectomy, radiation therapy, cryoablation therapy, n = 133) were included in the survival analysis (**Supplementary Table 5**). Proteins detected in at least 50% of samples (n = 164 including replicates, 1,936 proteins) were included. Primary outcome was time to biochemical recurrence (BCR), defined to be a sPSA rise of > 2 ng/mL above nadir for patients who underwent radiation therapy or cryoablation therapy, whereas BCR was defined with a sPSA ≥ 0.2 ng/mL post-treatment followed by a second confirmatory rising sPSA for patients received radical prostatectomy^26^. R packages survival (v.3.7-0) and survminer (v.0.4.9) were used for computing survival models. Univariate Cox proportional hazards regression model was applied to test the assumption of independent effects on association with outcome for each clinical variable (age, SIRE, cohort). A multivariable Cox proportional hazards regression model was applied to assess the association between outcome and multiple covariates, including log_2_ protein abundance treated as continuous variable, SIRE and cohort treated as categorical variables. An interaction term between protein abundance and SIRE was added to the model to assess interaction effects on outcome. The Wald-test p-value for each protein was corrected for multiple testing as described above^54^.

### Visualization and Statistical Analysis

The statistical tests used are indicated in the relevant figure legends or appropriate Methods section. Unless stated otherwise, statistical analyses and plotting were performed using R (v.4.3.0) and RStudio (version 2023.03.1+446). Data were visualized using R packages BoutrosLab.plotting.general^66^ (v.7.1.0), ggplot2 (v.3.5.1), ggbeeswarm (v.0.7.2), ggpubr (v.0.6.0), cowplot (v.1.1.3), ComplexHeatmap (v.2.16.0), ggrepel (v.0.9.6), circlize (v.4.16), gridExtra (v..2.3), corrplot (v.0.94), ggforitfy (v.0.4.17), gtsummary (v.2.0.2), and tidyverse (v.2.0.0).

## Acknowledgements

The authors thank all members of the Kislinger & Boutros Labs for their continued support and assistance.

## Data Availability

Raw mass spectrometry data and the generated spectral libraries are publicly available on MassIVE database with MassIVE ID: MSV000098127 and FTP link: ftp://MSV000098127@massive-ftp.ucsd.edu. Data supported corresponding results of the paper is included in the supplementary files.

## Author Contributions

A.H., A.K., B.P.M, J.O.N, and O.J.S contributed to data acquisition. A.H. and A.K. acquired the mass spectrometry data. A.H., A.K., J.A., and Z.Q. analyzed the data. J.O.N., P.C.B., and T.K. supervised the study. A.H., A.K., J.A., Z.Q., T.K., and P.C.B. wrote the manuscript, which all other authors edited and approved.

## References

1. Decramer, S. et al. Urine in clinical proteomics. Mol Cell Proteomics 7, 1850–1862 (2008).

2. Khoo, A. et al. Proteomic discovery of non-invasive biomarkers of localized prostate cancer using mass spectrometry. Nat Rev Urol 18, 707–724 (2021).

3. Ploussard, G. & de la Taille, A. Urine biomarkers in prostate cancer. Nat Rev Urol 7, 101–109 (2010).

4. Koo, K. M., Mainwaring, P. N., Tomlins, S. A. & Trau, M. Merging new-age biomarkers and nanodiagnostics for precision prostate cancer management. Nat Rev Urol 16, 302–317 (2019).

5. Tomlins, S. A. et al. Urine TMPRSS2:ERG Plus PCA3 for Individualized Prostate Cancer Risk Assessment. Eur Urol 70, 45–53 (2016).

6. Khoo, A. et al. Prostate cancer reshapes the secreted and extracellular vesicle urinary proteomes. Nat Commun 15, 5069 (2024).

7. Wilson, P. B., Winter, I. P. & Burdin, J. Differences in urine creatinine and osmolality between black and white Americans after accounting for age, moisture intake, urine volume, and socioeconomic status. PLoS One 19, e0304803 (2024).

8. Saka, A. H. et al. Cancer statistics for African American and Black people, 2025. CA Cancer J Clin (2025) doi:10.3322/caac.21874.

9. Delgado, C. Black men have higher rates of late stage prostate cancer, audit shows. BMJ 388, r50 (2025).

10. Garraway, I. P. et al. Prostate Cancer Foundation Screening Guidelines for Black Men in the United States. NEJM Evidence 3, (2024).

11. Klebaner, D. et al. Association of Health-Care System With Prostate Cancer-Specific Mortality in African American and Non-Hispanic White Men. JNCI: Journal of the National Cancer Institute 113, 1343–1351 (2021).

12. Dess, R. T. et al. Association of Black Race With Prostate Cancer–Specific and Other-Cause Mortality. JAMA Oncol 5, 975 (2019).

13. Jaratlerdsiri, W. et al. African-specific molecular taxonomy of prostate cancer. Nature 609, 552–559 (2022).

14. Yamaguchi, T. N. et al. The Germline and Somatic Origins of Prostate Cancer Heterogeneity. Cancer Discov 15, 988–1017 (2025).

15. Tosoian, J. J. et al. Prevalence and Prognostic Significance of PTEN Loss in African-American and European-American Men Undergoing Radical Prostatectomy. Eur Urol 71, 697–700 (2017).

16. Berchuck, J. E. et al. The Prostate Cancer Androgen Receptor Cistrome in African American Men Associates with Upregulation of Lipid Metabolism and Immune Response. Cancer Research 82, 2848–2859 (2022).

17. Mahal, B. A. et al. Prostate Cancer Genomic-risk Differences Between African-American and White Men Across Gleason Scores. European Urology 75, 1038– 1040 (2019).

18. Moul, J. W. et al. Prostate-specific antigen values at the time of prostate cancer diagnosis in African-American men. JAMA 274, 1277–1281 (1995).

19. Lai, S.-M., Keighley, J., Garimella, S., Enko, M. & Parker, W. P. Variations in Age-Adjusted Prostate Cancer Incidence Rates by Race and Ethnicity After Changes in Prostate-Specific Antigen Screening Recommendation. JAMA Netw Open 5, e2240657 (2022).

20. Li, C. H., Haider, S. & Boutros, P. C. Age influences on the molecular presentation of tumours. Nat Commun 13, 208 (2022).

21. Epstein, J. I. et al. The 2014 International Society of Urological Pathology (ISUP) Consensus Conference on Gleason Grading of Prostatic Carcinoma: Definition of Grading Patterns and Proposal for a New Grading System. Am J Surg Pathol 40, 244–252 (2016).

22. Yuan, J. et al. Integrative comparison of the genomic and transcriptomic landscape between prostate cancer patients of predominantly African or European genetic ancestry. PLoS Genet 16, e1008641 (2020).

23. Hänzelmann, S., Castelo, R. & Guinney, J. GSVA: gene set variation analysis for microarray and RNA-Seq data. BMC Bioinformatics 14, 7 (2013).

24. Subramanian, A. et al. Gene set enrichment analysis: A knowledge-based approach for interpreting genome-wide expression profiles. Proc. Natl. Acad. Sci. U.S.A. 102, 15545–15550 (2005).

25. Wang, Y. et al. Differentially expressed glycoproteins in pre– and post-digital rectal examination urine samples for detecting aggressive prostate cancer. Proteomics 23, e2200023 (2023).

26. Morgan, T. M. et al. Salvage Therapy for Prostate Cancer: AUA/ASTRO/SUO Guideline Part I: Introduction and Treatment Decision-Making at the Time of Suspected Biochemical Recurrence after Radical Prostatectomy. J Urol 211, 509– 517 (2024).

27. Tosoian, J. J. et al. Development and Validation of an 18-Gene Urine Test for High-Grade Prostate Cancer. JAMA Oncol 10, 726 (2024).

28. Kim, Y. et al. Targeted proteomics identifies liquid-biopsy signatures for extracapsular prostate cancer. Nat Commun 7, 11906 (2016).

29. Soh, P. X. Y. et al. Prostate cancer genetic risk and associated aggressive disease in men of African ancestry. Nat Commun 14, 8037 (2023).

30. Arenas-Gallo, C. et al. Race and prostate cancer: genomic landscape. Nat Rev Urol 19, 547–561 (2022).

31. Rayford, W. et al. Comparative analysis of 1152 African-American and European-American men with prostate cancer identifies distinct genomic and immunological differences. Commun Biol 4, 670 (2021).

32. Awasthi, S. et al. Comparative Genomics Reveals Distinct Immune-oncologic Pathways in African American Men with Prostate Cancer. Clinical Cancer Research 27, 320–329 (2021).

33. Weiner, A. B. et al. Plasma cells are enriched in localized prostate cancer in Black men and are associated with improved outcomes. Nat Commun 12, 935 (2021).

34. Wallace, T. A. et al. Tumor Immunobiological Differences in Prostate Cancer between African-American and European-American Men. Cancer Research 68, 927–936 (2008).

35. Sartor, O. et al. Survival of African-American and Caucasian men after sipuleucel-T immunotherapy: outcomes from the PROCEED registry. Prostate Cancer Prostatic Dis 23, 517–526 (2020).

36. Kiely, M. & Ambs, S. Immune Inflammation Pathways as Therapeutic Targets to Reduce Lethal Prostate Cancer in African American Men. Cancers 13, 2874 (2021).

37. Feibus, A. H. et al. Clinical Use of PCA3 and TMPRSS2:ERG Urinary Biomarkers in African-American Men Undergoing Prostate Biopsy. Journal of Urology 196, 1053– 1060 (2016).

38. Yuan, J. et al. Integrated Analysis of Genetic Ancestry and Genomic Alterations across Cancers. Cancer Cell 34, 549–560.e9 (2018).

39. Sucheston, L. E., et al. Genetic Ancestry, Self-Reported Race and Ethnicity in African Americans and European Americans in the PCaP Cohort. PLoS ONE 7, e30950 (2012).

40. Banda, Y. et al. Characterizing Race/Ethnicity and Genetic Ancestry for 100,000 Subjects in the Genetic Epidemiology Research on Adult Health and Aging (GERA) Cohort. Genetics 200, 1285–1295 (2015).

41. Ha, A. et al. Comprehensive Prostate Fluid-Based Spectral Libraries for Enhanced Protein Detection in Urine. J. Proteome Res. 23, 1768–1778 (2024).

42. Berger, S. T. et al. MStern Blotting-High Throughput Polyvinylidene Fluoride (PVDF) Membrane-Based Proteomic Sample Preparation for 96-Well Plates. Mol Cell Proteomics 14, 2814–2823 (2015).

43. Escher, C. et al. Using iRT, a normalized retention time for more targeted measurement of peptides. Proteomics 12, 1111–1121 (2012).

44. Searle, B. C. et al. Chromatogram libraries improve peptide detection and quantification by data independent acquisition mass spectrometry. Nat Commun 9, 5128 (2018).

45. MacLean, B., et al. Skyline: an open source document editor for creating and analyzing targeted proteomics experiments. Bioinformatics 26, 966–968 (2010).

46. Yu, F. et al. Analysis of DIA proteomics data using MSFragger-DIA and FragPipe computational platform. Nat Commun 14, 4154 (2023).

47. Kong, A. T., Leprevost, F. V., Avtonomov, D. M., Mellacheruvu, D. & Nesvizhskii, A. I. MSFragger: ultrafast and comprehensive peptide identification in mass spectrometry-based proteomics. Nat Methods 14, 513–520 (2017).

48. da Veiga Leprevost, F., et al. Philosopher: a versatile toolkit for shotgun proteomics data analysis. Nat Methods 17, 869–870 (2020).

49. Demichev, V., Messner, C. B., Vernardis, S. I., Lilley, K. S. & Ralser, M. DIA-NN: neural networks and interference correction enable deep proteome coverage in high throughput. Nat Methods 17, 41–44 (2020).

50. Yu, F., Haynes, S. E. & Nesvizhskii, A. I. IonQuant Enables Accurate and Sensitive Label-Free Quantification With FDR-Controlled Match-Between-Runs. Mol Cell Proteomics 20, 100077 (2021).

51. Yang, K. L. et al. MSBooster: improving peptide identification rates using deep learning-based features. Nat Commun 14, 4539 (2023).

52. Ritchie, M. E. et al. limma powers differential expression analyses for RNA-sequencing and microarray studies. Nucleic Acids Research 43, e47–e47 (2015).

53. Stekhoven, D. J. & Bühlmann, P. MissForest—non-parametric missing value imputation for mixed-type data. Bioinformatics 28, 112–118 (2012).

54. Benjamini, Y. & Hochberg, Y. Controlling the False Discovery Rate: A Practical and Powerful Approach to Multiple Testing. Journal of the Royal Statistical Society Series B: Statistical Methodology 57, 289–300 (1995).

55. Venables, W. N. & Ripley, B. D. *Modern Applied Statistics with S*. (Springer New York, New York, NY, 2002). doi:10.1007/978-0-387-21706-2.

56. Goldman, M. J. et al. Visualizing and interpreting cancer genomics data via the Xena platform. Nat Biotechnol 38, 675–678 (2020).

57. Price, A. L. et al. Principal components analysis corrects for stratification in genome-wide association studies. Nat Genet 38, 904–909 (2006).

58. Uhlén, M. et al. Proteomics. Tissue-based map of the human proteome. Science 347, 1260419 (2015).

59. Karlsson, M. et al. A single-cell type transcriptomics map of human tissues. Sci Adv 7, eabh2169 (2021).

60. Uhlen, M. et al. A genome-wide transcriptomic analysis of protein-coding genes in human blood cells. Science 366, eaax9198 (2019).

61. Liberzon, A. et al. The Molecular Signatures Database Hallmark Gene Set Collection. Cell Systems 1, 417–425 (2015).

62. Milacic, M. et al. The Reactome Pathway Knowledgebase 2024. Nucleic Acids Research 52, D672–D678 (2024).

63. The Gene Ontology Consortium et al. The Gene Ontology knowledgebase in 2023. GENETICS 224, iyad031 (2023).

64. Shannon, P. et al. Cytoscape: A Software Environment for Integrated Models of Biomolecular Interaction Networks. Genome Res. 13, 2498–2504 (2003).

65. D’Amico, A. V. et al. Biochemical outcome after radical prostatectomy, external beam radiation therapy, or interstitial radiation therapy for clinically localized prostate cancer. JAMA 280, 969–974 (1998).

66. P’ng, C. et al. BPG: Seamless, automated and interactive visualization of scientific data. BMC Bioinformatics 20, 42 (2019).

