## Supplementary Information for "Ancestry-Dependent Immunologic and Prognostic Effects Characterize the Prostate Cancer Urinary Proteome"

* equal contributions

**Supplementary Figures**

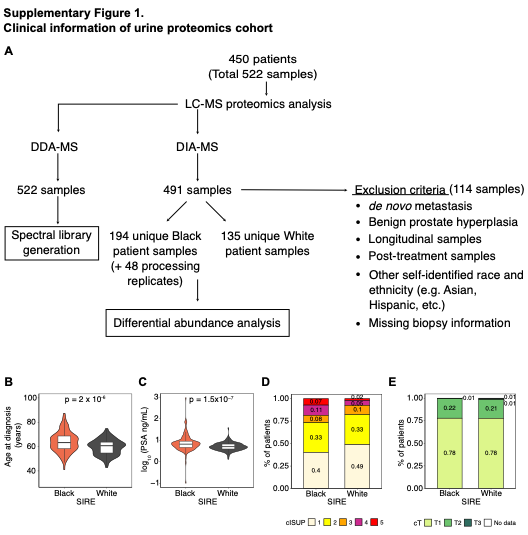

**Supplementary Figure 1. Clinical information of urine proteomics cohort.** (A) Clinical patient samples inclusion flowchart. (B - C) Violin plots and boxplots showing the distributions of (B) age and (C) log_10_ serum PSA (sPSA) level at diagnosis between Black and White patients. P-values from Wilcoxon’s test are shown. (D - E) Stacked bar plots of the proportion of patients’ clinical stage (D) cISUP grade group, and (E) clinical T category, between Black and White patients.

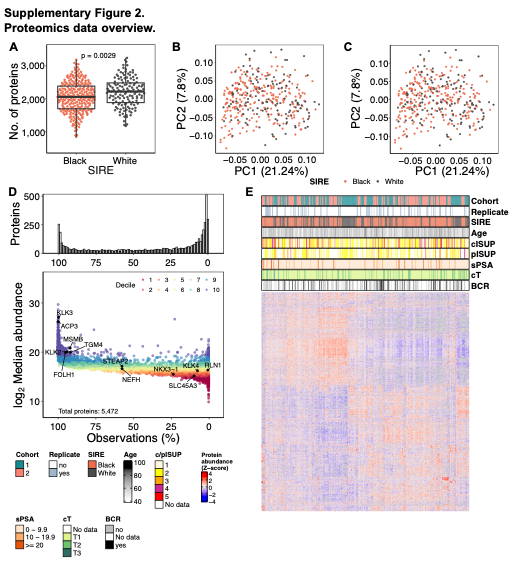

**Supplementary Figure 2. Overview of proteomics data.** (A) Number of proteins detected per sample across Black and White patients. P-value from Mann-Whitney U test is shown. (B - C) Principal component analysis (PCA) dimensions 1 and 2 of the protein abundance (B) prior to and (C) after data normalization and batch correction. (D) Overall protein detection rates of the processed samples (n = 377). Top: histogram of protein detection rate across the cohorts. Bottom: distribution of median log_2_ protein abundance coloured by decile. (E) Heatmap of proteins included in subsequent multivariable analysis (n = 1,782) with protein abundance scaled as z-score.

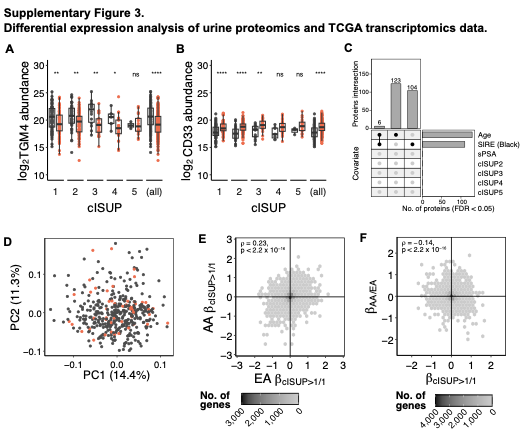

**Supplementary Figure 3. Differential abundance of urine proteomics and TCGA tissue transcriptomics data.** (A - B) Protein abundance of top ranking proteins (A) TGM4 and (B) CD33 across cISUP grade group between Black and White patients. (dark grey: White patients; red orange: Black patients). P-values from Mann-Whiteney U test comparing protein abundance within each cISUP grade group between SIRE are shown as asterisks (ns: not significant; *: p < 0.05; **: p < 0.01; ***: p < 0.001). (C) Upset plot of significantly associated proteins (FDR < 0.05) for each clinical covariate against baseline in urine proteomics multivariable analysis (SIRE:White, cISUP: 1). (D) PCA of TCGA-PRAD transcriptomics data stratified by genetically inferred ancestry (dark grey: EA patients; red orange: AA patients). (E - F) Spearman’s correlations of log_2_RSEM associations with (E) disease stages (cISUP>1/cISUP1) between AA and EA patients from stratified analyses within ancestry, (F) ancestry and disease stage from multivariable analysis not stratified.

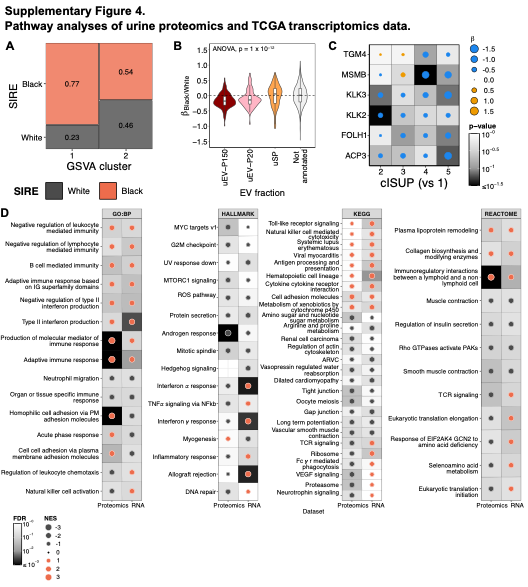

**Supplementary Figure 4. Annotations and pathway analyses of -omics data.** (A) Mosaic plot of the proportion of samples in the clusters separated by the second-degree column dendrogram of the GSVA unsupervised hierarchical clustering. (B) Violin plot and boxplot of protein association with SIRE (β) annotated by urinary extracellular vesicle (uEV) fractions (Khoo *et al*. Nat Comms 2024). (C) Regression coefficients of prostate-derived proteins for cISUP 2 – 5 against cISUP 1 from multivariable analysis adjusting for age, sPSA and SIRE. Dot size represents magnitude of effect size, colour represents directionality (blue: negative associated; orange: positively associated), shading represents the p-value from t-statistics of linear regression. (D) Dotplot of the pre-ranked GSEA pathway enrichment analysis from both urine proteomics and tissue transcriptomics multivariable linear regression analyses. Pathways from GO:BP, HALLMARK, KEGG, and REACTOME databases with FDR < 0.25 for both or either -omics datasets are shown. Dot size represents the absolute NES values, colour represents direction of enrichment (red orange: positive and enriched in AA; dark grey: negative and enriched in EA), and background shaking represents statistical significance.

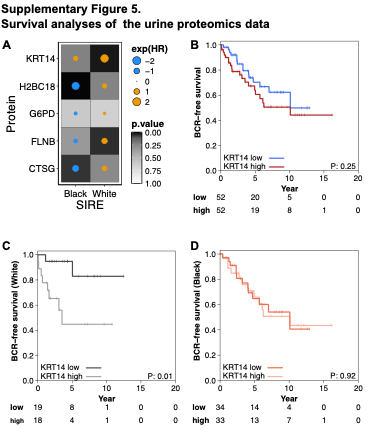

**Supplementary Figure 5. Survival analyses of the urine proteomics data.** (A) Dot map showing the protein abundance association with BCR of the top 5 ranking proteins in independent analysis stratified within Black and White patients. Dot size shows the absolute regression coefficient of the protein abundance association with BCR, colour represents the directionality of regression (blue: better prognosis; orange: worse prognosis), background shading represents the significance of association from the Wald statistics. (B) KM plot of intermediate patients stratified by KRT14 abundance (median dichotomized) only. (C – D) KM plot of intermediate patients stratified by KRT14 abundance (median dichotomized), within (C) White patients and (D) Black patients only. P-values from the model log-rank test are shown on the KM plots.

**Supplementary Tables**

**Supplementary Table 1:** Overall patient clinical characteristics.

**Supplementary Table 2:** Urine proteomics data used for differential expression analysis.

Protein abundance (n = 1,785) data for the included patient samples (n = 377) after normalization and filtering.

**Supplementary Table 3:** Urine proteomics differential expression analyses results.

Univariable and multivariable analysis results of protein abundance association with self-identified race and ethnicity. Gene set enrichment analysis results were calculated based on the regression coefficients of Black/White in the multivariable analysis adjusting for age, sPSA, cISUP, and cohort.

**Supplementary Table 4:** TCGA-PRAD tissue transcriptomics differential expression analysis results.

**Supplementary Table 5:** Biochemical recurrence-free survival analysis results.

Patients of D’Amico intermediate risk group (n = 135) are included for the analysis. Overall clinical Cox-proportional hazards univariable and multivariable model results are reported.
